# Analysis of MPXV RNA-seq Data Reveals Lack of Evidence of APOBEC3-mediated RNA Editing

**DOI:** 10.1101/2025.07.21.665962

**Authors:** Alisa O. Lyskova, Ruslan Kh. Abasov, Anna Pavlova, Evgenii V. Matveev, Alexandra Madorskaya, Fedor M. Kazanov, Daria V. Garshina, Anna E. Smolnikova, Gennady V. Ponomarev, Elena I. Sharova, Dmitry N. Ivankov, Ogun Adebali, Mikhail S. Gelfand, Marat D. Kazanov

**Author notes:** equal contribution (Alisa O. Lyskova, Ruslan Kh. Abasov).

## Abstract

The 2022 outbreak of monkeypox virus (MPXV), a double-stranded DNA virus, is remarkable for an unusually high number of single-nucleotide substitutions compared to earlier strains, with a strong bias toward C→T and G→A transitions consistent with the APOBEC3 cytidine deaminase activity. While APOBEC3-induced mutagenesis is well documented at the DNA level, its potential impact on MPXV RNA transcripts remains unclear. To assess whether APOBEC3 enzymes act on MPXV RNA, we analyzed RNA-seq data from infected samples. The enrichment of APOBEC-signature substitutions among high-frequency mismatched positions led us to consider two possibilities: RNA editing at hotspots or fixed DNA mutations. Multiple lines of evidence support the conclusion that these substitutions arise from DNA-level mutagenesis rather than RNA editing. These include a substantial number of G→A substitutions remaining after normalization by gene strand direction, a largely neutral impact of substitutions on protein-coding sequences, the lack of positional correlation with transcriptional features or RNA secondary structure typically associated with APOBEC action hotspots, and an overlap with known genomic mutations in MPXV strains. Analysis of the nucleotide context of observed substitutions indicated that APOBEC3A or APOBEC3B were likely drivers of DNA-level mutagenesis.

**Importance:** The 2022 monkeypox virus (MPXV) outbreak showed an unusually high number of mutations thought to result from human antiviral enzymes of the APOBEC3 family. While such mutations have been clearly documented in the viral DNA, whether APOBEC3 also edits viral messenger RNA molecules remained unclear. In this study, we analyzed multiple publicly available MPXV RNA sequencing datasets to address this question. We found that the apparent APOBEC-like changes in RNA are best explained by fixed DNA mutations rather than active RNA editing. This finding helps clarify how MPXV evolves and adapts, suggesting that APOBEC3’s role in shaping the virus likely operates at the DNA level. Understanding where and how these mutations occur provides insight into the virus’s interaction with the human immune system and informs future studies on viral evolution and antiviral defenses.

## Introduction

Monkeypox disease, known for a multi-country outbreak in 2022, is caused by the monkeypox virus (MPXV), a double-stranded DNA virus belonging to the Orthopoxvirus genus in the Poxviridae family [1]. MPXV was first isolated in 1958 in Copenhagen from a Macaca fascicularis imported from Singapore [1]. The first human case of MPXV was documented in 1970 in the Congo Basin [2], demonstrating the virus’s capability for a zoonotic transmission from animals to humans. Since then, numerous outbreaks were reported across Eastern, Central, and Western Africa [3]. The symptoms of monkeypox are similar to those of smallpox, which also belongs to the Orthopoxvirus genus along with cowpox, camelpox, mousepox, and vaccinia viruses [4]. However, monkeypox typically has a lower mortality rate than smallpox [5]. Non-human primates are susceptible to MPXV and can transmit the virus during outbreaks. However, there is no clear evidence supporting long-term maintenance of the virus within primate populations [6]. The natural reservoir of MPXV is thought to be rodents and other small animals [7]. From 2002 to 2021, several cases of MPXV reported outside of Africa were associated with imports from endemic countries in West and Central Africa [8,9]. In 2022, a rapidly expanding multi-country outbreak affected individuals with no travel history to Africa, suggesting that MPXV may have evolved to enable human-to-human transmission [10]. Since the start of the outbreak, more than 90,000 cases were reported across over 110 countries, leading the World Health Organization (WHO) to declare it a global health emergency.

MPXV is phylogenetically divided into three distinct clades: Clade I, recently subdivided into Ia and Ib, represent the most virulent lineage with up to 10% mortality in humans and is predominantly transmitted in the Congo Basin; Clade IIa, initially a zoonotic strain with low mortality in West Africa, later adapted to the human-to-human transmission and caused an outbreak in Nigeria during 2017–2018; and Clade IIb, associated with the 2022 global outbreak, characterized by its ability to spread through human-to-human transmission [11,12]. The MPXV genome is approximately 197,000 base pairs long and contains about 200 genes [6]. A relatively large number of genes are involved in to the encoded replication machinery, as the entire viral lifecycle takes place in the cytoplasm of infected cells [13]. The MPXV genome includes two telomeres with identical sequences that are oriented in opposite directions, known as inverted terminal repeats (ITRs) [14]. Genes responsible for replication and morphogenesis, conserved among poxviruses, are clustered in the central region of the genome, while the genes specific to pathogenesis are located in the terminal regions [15]. Studies on the transcriptional program of MPXV categorized its genes into three temporal expression phases: early, intermediate, and late [16]. The estimated mutation rate for orthopoxviruses is 1–2 substitutions per genome per year [17], however, the extent of divergence in MPXV genomes from the 2022 outbreak, compared to the related 2018–2019 viruses, has reached approximately 50 single-nucleotide substitutions [12]. Most substitutions were C→T transitions or their complementary G→A counterparts, typically occurring in the TC (complementary GA) contexts, respectively [12]. This mutational signature, characteristic of APOBEC3 family cytidine deaminases, suggests that the accumulation of these mutations in the MPXV genome could indicate the virus’s evolved ability for human-to-human transmission [12,18,19].

Apolipoprotein B mRNA Editing Catalytic Polypeptide-like (APOBEC) is a family of enzymes in mammals that play a key role in the immune system, restricting viruses and retrotransposons [20]. Recently, enzymes of the APOBEC3 subfamily have received significant attention for their role in cancer mutagenesis and its potential involvement in the mutagenesis of the SARS-CoV-2 virus [21–23]. APOBEC enzymes bind to RNA or single-stranded DNA, catalyzing deamination of cytosine to uracil. In an RNA molecule, the altered nucleotide remains unchanged, as uridine is a normal RNA base and RNA is not subject to the uracil-removal repair pathways that operate on DNA. In a DNA molecule, uracil can be fixed as thymine in subsequent rounds of replication or, if converted by DNA glycosylases into an abasic site, it may be further repaired by translesion polymerases into either thymine or guanine [24]. It has been observed that there exist mutagenesis hotspots of APOBEC enzymes occurring within particular secondary structure patterns of RNA or single-stranded DNA [25]. Specifically, a cytosine located at the 3’ end of a hairpin loop would have a 10-fold higher mutation rate by APOBEC enzymes compared to cytosines in TC motifs outside secondary structure [26]. A similar effect was observed for APOBEC signature mutations in the MPXV genome, providing additional support for the involvement of APOBEC enzymes [27]. While mutations in MPXV genomes associated with the 2022 outbreak have been extensively studied, it remains unclear whether viral messenger RNAs are also affected by APOBEC enzymes.

In this study, we analyzed publicly available MPXV RNA-seq data for evidence of APOBEC mutagenesis in viral transcripts. Analysis of MPXV transcriptomes generated by RNA-sequencing can provide valuable insights into the extent and characteristics of APOBEC mutagenesis. Moreover, RNA-seq studies can generate significantly more statistical data than genome sequencing, as messenger RNAs are produced in multiple copies during gene expression, and each of these molecules can theoretically be targeted by APOBEC enzymes. Our results reveal a moderate number of APOBEC-signature substitutions in MPXV transcripts which, upon thorough analysis, appear to stem from DNA-level mutagenesis, hence limiting support for APOBEC-mediated RNA editing.

## Methods

RNA-seq datasets (Table S1) from the following projects were obtained from the SRA archive: PRJEB60728 (host: *Homo sapiens*), PRJNA906618 (host: *Homo sapiens*), PRJEB56841 (host: *Chlorocebus sabaeus*), PRJNA1183318 (host: *Macaca fascicularis*) along with DNA sequencing data from projects PRJNA845087 and PRJNA981509 (both host: *Homo sapiens*). Both RNA-seq and DNA-seq datasets were downloaded using the fastq-dump utility from the SRA Toolkit (version 3.1.1) [28][29]. The reference genome of the monkeypox virus (MPXV) (GCF_014621545.1 / ASM1462154v1), as well as host genomes (*Homo sapiens:* GRCh38.d1.v1, *Chlorocebus sabaeus:* GCF_015252025.1, *Macaca fascicularis:* GCA_037993035.1), were obtained from the NCBI RefSeq database [29][30].

RNA-seq data generated using long-read sequencing technologies were aligned to the reference genome using minimap2 (version 2.28-r1209) [30][31]. Short-read RNA-seq data were aligned using STAR (version 2.7.11b) [31][32], and genomic DNA short reads were mapped using BWA (version 2.2.1) [32] [33]. Alignment quality metrics were computed using mosdepth (version 0.3.9) [33][34] and the samtools stats command from samtools (version 1.20) [34][35]. Detection of positions with mismatches, representing either genomic variants or potential RNA editing sites, was performed using BCFtools (version 1.21, the exact command-line parameters are provided in Table S21) [34][35]. Mismatch positions were filtered using a substitution frequency threshold of 1% for datasets generated with short-read sequencing, which is approximately ten times higher than the reported sequencing error rate [35][36]. For datasets based on long-read sequencing, a more conservative threshold of 10% was applied due to the higher intrinsic error rate of this technology. Read counts were generated using the HTSeq package [36], and transcript abundance was subsequently quantified in TPM units using the edgeR package [37].

Secondary structure around the identified mismatch positions was predicted using RNAfold [38] and further analyzed with the RNAsselem package (version 1.3) [39][38]. Inverted repeats were identified using Palindrome Analyzer [40][39]. The genomic map was visualized using the circlize package in R. All other plots were generated using ggplot2 (version 3.5.1) and matplotlib (version 3.9.0).

To model the sequence specificity of APOBEC family enzymes (APOBEC3A, APOBEC3B, APOBEC3F, and APOBEC3G), we constructed position weight matrices (PWMs) based on published data reporting experimentally confirmed mutation sites and their sequence contexts [41–43][40–42]. For each enzyme, we compiled nucleotide sequences flanking known editing sites from the literature and counted the frequency of each nucleotide (A, C, G, T) at each position within a defined 5-nucleotide window (–3, –2, –1, 0, 1) surrounding the mutation site, where position 0 corresponds to the mutated nucleotide. These nucleotide counts were assembled into a matrix, with each row corresponding to a nucleotide and each column to a relative position. The counts in each column were normalized to sum to 1, and the resulting frequencies were log-transformed to generate the final PWM. To score a given sequence position, we summed the logarithmic weights corresponding to the observed nucleotide at each position in the matrix. Additionally, PWMs specifically designed to distinguish APOBEC3A from APOBEC3B were adopted from [44][43], based on the differential preference for YTCA versus RTCA sequence motifs (the mutated C is underlined).

To assess where APOBEC-signature substitutions were enriched within each sample, we modeled the number of such substitutions using a beta-binomial distribution, which accounts for overdispersion relative to a simple binomial model. Here, an APOBEC-signature substitution was defined as a C→T substitution occurring in a TC context or the complementary G→A substitution in a GA context, consistent with the known sequence preference of APOBEC3 enzymes. Let ***n*** be the total number of detected substitutions in the sample, *k* the number of APOBEC-signature substitutions, and *π* the unknown underlying probability that a substitution belongs to the APOBEC-signature class. We assumed that *π* follows a beta prior: *π* ~ Beta(α, β), where the parameters α and β were estimated from the background (non-APOBEC-signature) substitutions across the sample. Under this model, the likelihood of observing *k* APOBEC-signature substitutions out of n total substitutions follows the beta-binomial distribution:

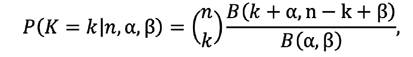

where *B*(,) denotes the beta function. Using the estimated parameters 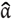 and 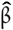, we computed the p-value for enrichment as the probability of observing *k* or more APOBEC-signature substitutions under the fitted beta-binomial model:

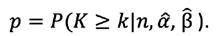

This test quantifies whether the observed proportion of APOBEC-signature substitutions exceeds the expectation defined by the background substitution process.

## Results

### APOBEC-signature substitutions in MPXV transcripts are observed at a limited number of positions, where they occur at high frequency

To identify RNA editing events in MPXV transcripts, we comprehensively analyzed available RNA-seq data from infected hosts, including two human [45,46], and two macaque studies [47,48] (hereafter referred to as RNAseq:human1–2 and RNAseq:monkey1–2), comprising a total of 47 samples. To ensure accurate detection of potential RNA editing events, we mapped sequenced reads to the MPXV genome using three complementary approaches: (i) direct mapping to the MPXV genome, (ii) mapping to the combined host and MPXV genome, and (iii) initial mapping to the host genome, followed by mapping of the unmapped reads to the MPXV genome. Following read mapping, we identified positions with mismatches using BCFtools allowing for highly sensitive detection of substitutions. To achieve maximum specificity, we set the threshold for substitution frequency as low as possible, but substantially higher than the probability of sequencing errors (see Methods). We applied this method to three alignments generated for each RNA-seq study and derived the consensus positions (Fig. 1a). Subsequent analysis of the detected transcript positions revealed similar results across the examined projects (Figures S1, S2), with Figure 1 illustrating a representative example using a single sample ERR11030177 from project [45]. In this sample, more than 569 positions with potential RNA editing events were identified using the BCFtools method. We analyzed the fractions of substitution types at the detected positions that could be attributed to APOBEC-signature substitutions, specifically C→T or complementary G→A changes in the TC/GA context. Alternatively, we considered the same substitutions occurring outside of this context as part of the extended APOBEC signature [49]. We observed a moderate enrichment of C→T and complementary G→A substitutions (Fig. 1b) and analyzed their two-nucleotide context, revealing that a significant fraction of these substitutions (p = 7.09 × 10^−8^, see Methods) had occurred within the APOBEC TC/GA motif (Fig. 1b, 1c). However, the analysis of distribution of substitution frequencies (SF) at detected positions (Fig. 1d) revealed that most APOBEC-signature positions have SF greater than 50% (p = 3.75 × 10^−9^), whereas positions with medium and low SF show no enrichment (p = 0.33) of APOBEC-signature substitutions (Fig. 1e). We assumed that the high frequency of APOBEC-signature substitutions at specific transcript positions could result from either pre-existing DNA-level substitutions or RNA editing hotspots within transcripts. To distinguish between these alternatives, we conducted an additional analysis related to the identified positions. For subsequent analysis, we first filtered substitution positions by substitution frequency (SF > 0.5) to retain the range enriched for APOBEC-signature substitutions and then compiled a non-redundant list of substitution positions observed across all samples in each study, indicating the number of samples in which each position occurred (Supplemental file 1).

**Figure 1.**
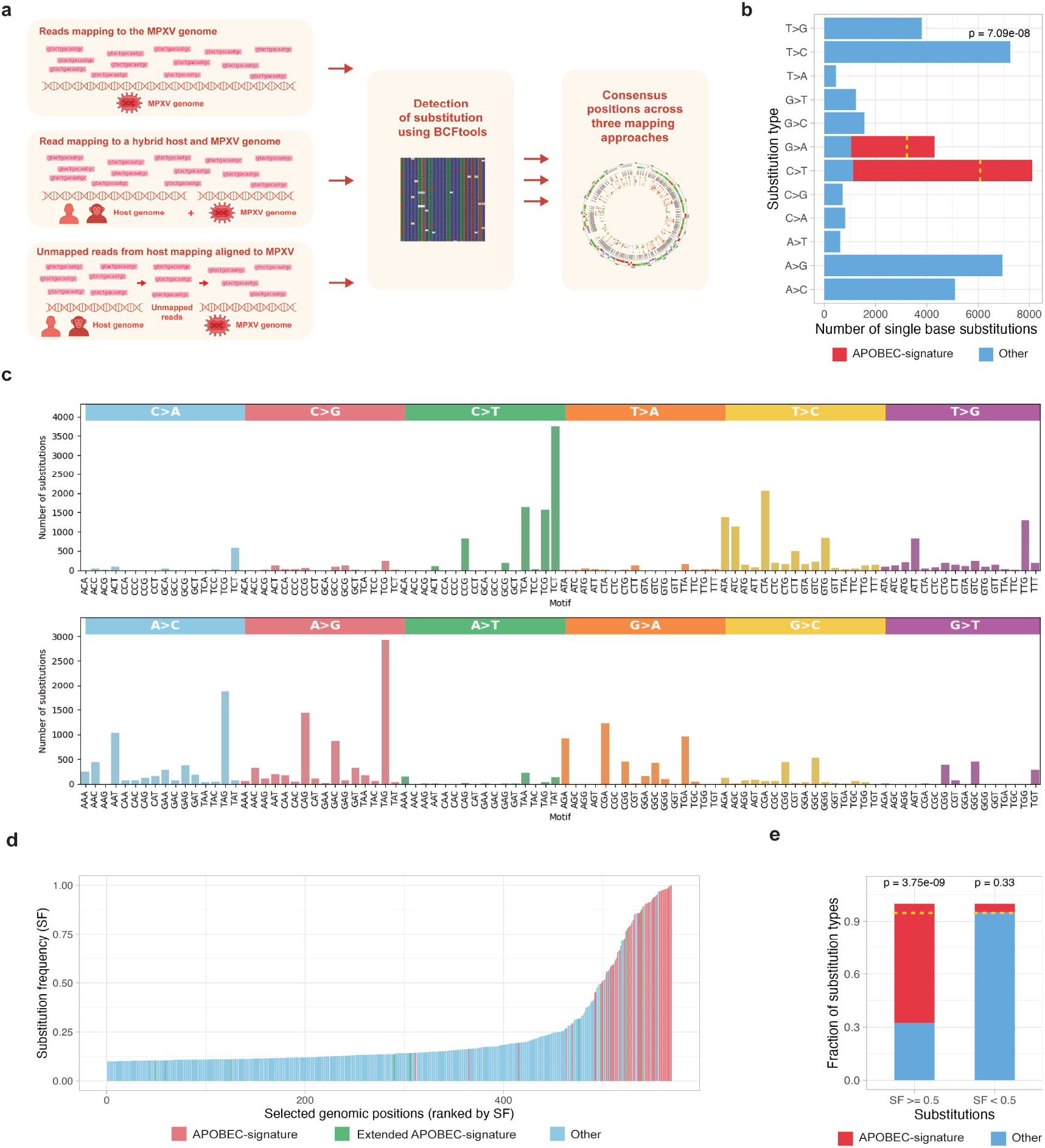
**(a)** Scheme of the pipeline for detection of variant-like positions in RNA-seq data, which may reflect either genomic variants or RNA editing events. **(b)** Distribution of single-base substitution types detected in sample. Bars correspond to observed counts for each substitution class, with APOBEC-signature substitutions highlighted in red. The dotted yellow line marks the expected number of APOBEC-signature substitutions estimated from the beta-binomial model. **(c)** Distribution of single-nucleotide substitutions in MPXV transcripts across all trinucleotide sequence contexts, grouped by substitution type. Each bar represents the number of observed substitutions occurring in a specific 3-nt motif, with colored panels indicating major substitution classes. **(d)** Substitution frequencies of all detected mismatch positions in sample, sorted by increasing frequency. Each bar represents one genomic position, ordered on the x-axis by increasing substitution frequency (y-axis). Colors indicate the substitution class: APOBEC-signature C→T/G→A substitutions in the TC/GA motif (red), the same substitutions outside this motif (extended APOBEC signature; green), and all other substitution types (blue). (>0.5). **(e)** Enrichment of APOBEC-signature substitutions among high-frequency mismatches in sample. Bars show the proportions of APOBEC-signature (red) and other (blue) substitutions within two substitution frequency (SF) categories: high-frequency (SF ≥ 0.5) and low-frequency (0.1 ≤ SF < 0.5) sites. Horizontal dashed lines indicate the expected fraction of APOBEC-signature substitutions under the beta-binomial model. P-values above bars quantify the deviation from the expected substitution fractions. High-frequency mismatches show a significant enrichment of APOBEC-signature substitutions (left bar), whereas low-frequency mismatches do not show deviation from expectation (right bar).

### Gene-focused analysis at positions of APOBEC-signature substitutions provides evidence that a significant fraction of substitutions occurred at the DNA level

When APOBEC-signature RNA editing occurs in a transcript, non-stranded RNA sequencing can generate both direct and complementary reads, yielding both C→T and G→A substitutions in the TC/GA context. However, considering gene direction, we can assess whether suspected RNA editing leads to C→T substitutions in genes on the plus strand and G→A substitutions in genes on the minus strand of the double-stranded MPXV DNA virus. We normalized all substitutions according to the gene strand direction, preserving those in plus-strand gene regions and converting those in minus-strand gene regions to their complementary counterparts. Following this normalization, the proportion of G→A substitutions in human dataset decreased from 57.9% to 45.6% (non-human primates: from 55.4% to 53.6%), but still remained substantial (Fig. 2a, Fig. S3a). We conclude that a likely explanation for this is that a significant fraction of substitutions occurred not in RNA transcripts, but at the DNA level before transcription.

**Figure 2.**
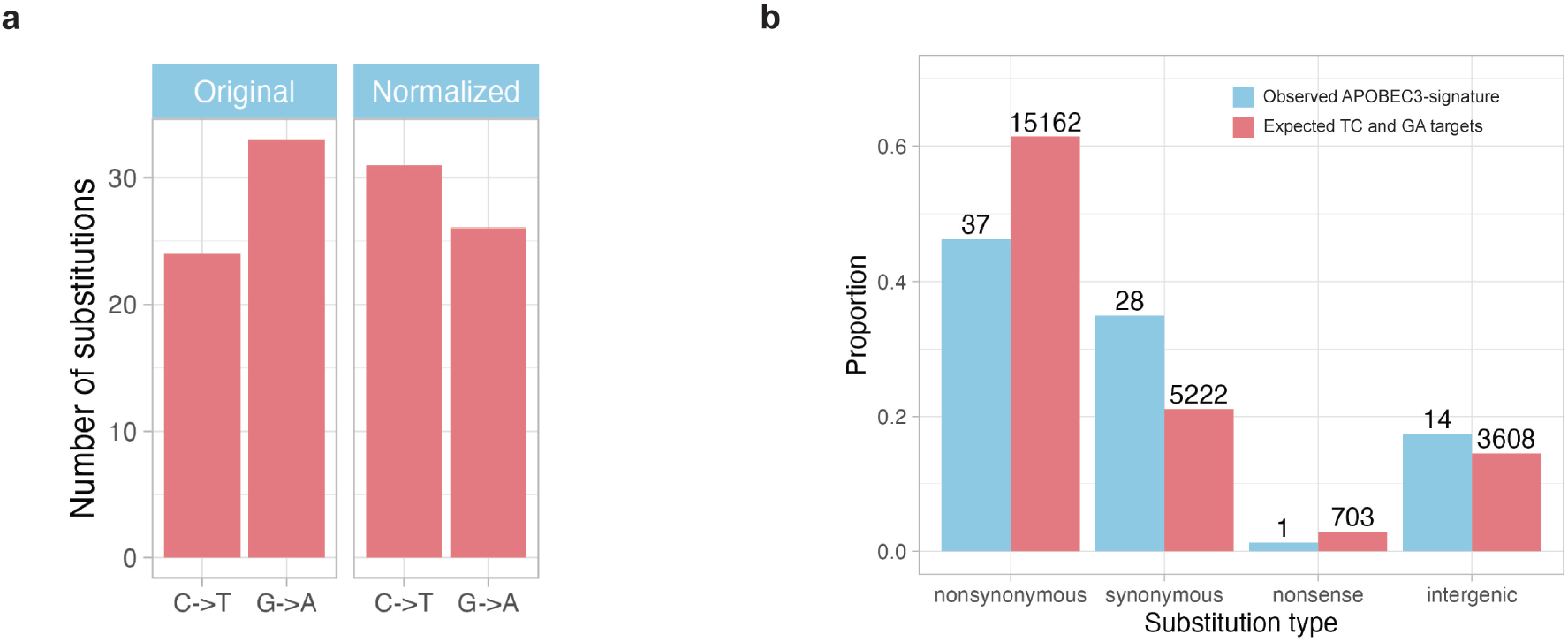
**(a)** Proportion of G→A substitutions with substitution frequency (SF) > 0.5 identified in the RNA-seq:human1-2 dataset before and after strand-based normalization of APOBEC-signature substitutions. **(b)** Distribution of APOBEC-signature substitutions identified in the RNA-seq:human1–2 dataset across functional mutation categories, including synonymous, non-synonymous, nonsense, and intergenic substitutions.

**Figure 3.**
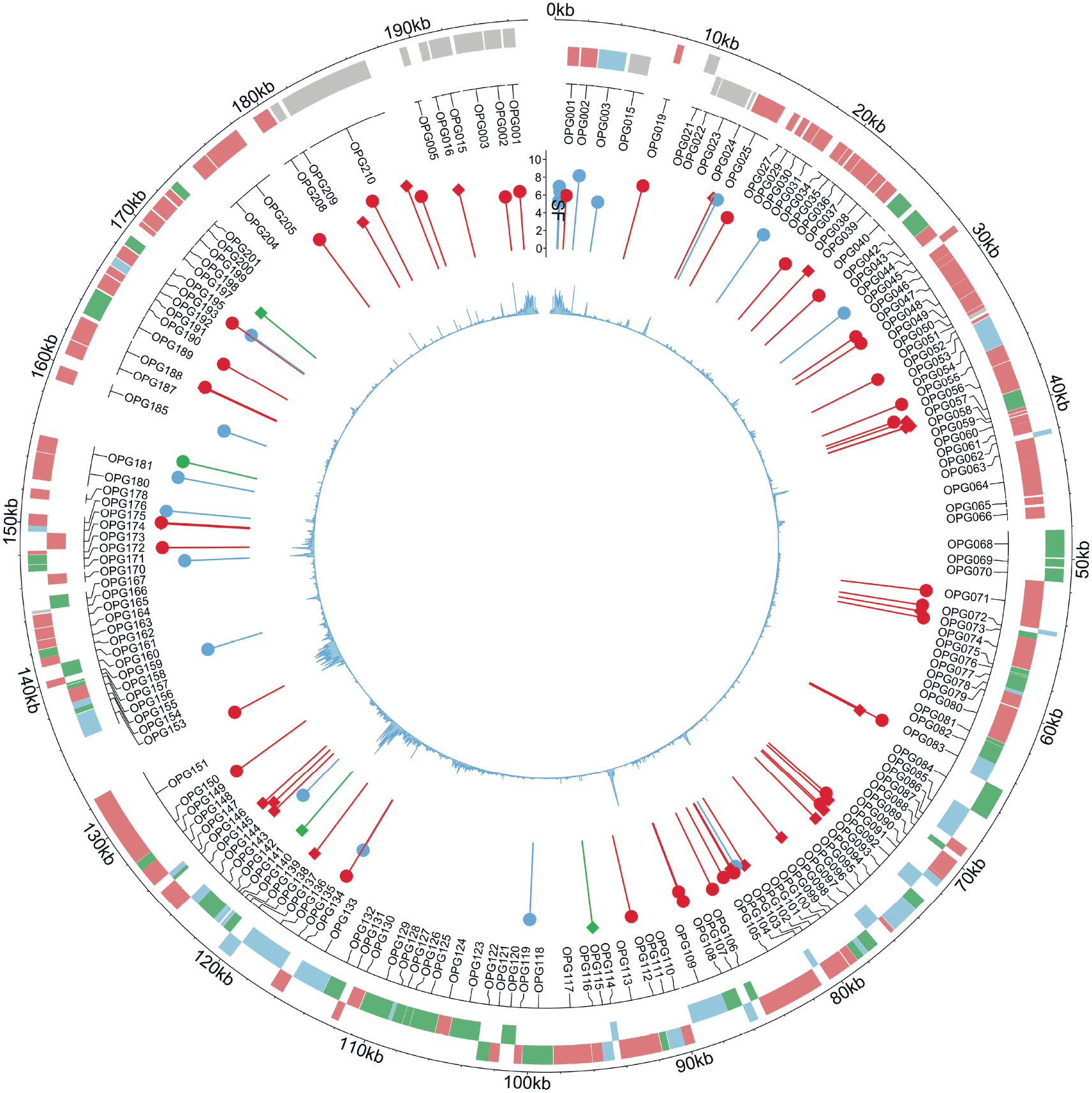
Genomic map of identified substitution positions in the MPXV genome from two human-derived projects. The outermost track shows annotated viral genes, color-coded by expression phase: early (green), middle (red), and late (blue). The next layer inward displays the positions of identified substitutions. The height of each pin reflects the substitution frequency (SF). Pin color indicates substitution type: APOBEC-signature (red), extended APOBEC-signature (green), and other substitutions (blue). The pin head shape denotes strand orientation: a circle indicates a C→T substitution on the gene strand, while a square indicates the same substitution on the complementary strand. The innermost track displays the mean coverage across the genome.

Additional support for this hypothesis could come from estimating the effect of APOBEC-signature substitutions on the encoded amino acid sequence of transcripts. If these substitutions occur at the DNA level, deleterious substitutions may severely disturb the viral life cycle. Hence, most observed substitutions should be neutral or at worst mildly deleterious. We classified APOBEC-signature substitutions based on their impact on gene function and observed a depletion of non-synonymous substitutions (binomial test: human, p = 4.23 × 10^−3^; non-human primates, p = 8.01 × 10^−3^), which are often deleterious, along with an enrichment of synonymous ones (binomial test: human, p = 2.98 × 10^−3^; non-human primates, p = 1.9 × 10^−4^), which are generally considered neutral (Fig. 2b, Fig. S3b).

### APOBEC-signature substitutions do not correlate with MPXV genomic features associated with transcription

To further distinguish between two hypotheses about the observed APOBEC signature substitutions – whether they result from RNA editing at transcript hotspots or from DNA-level substitutions – we analyzed the correlation between the identified genomic positions and known transcription-associated genomic features. If RNA editing predominates, we may expect the detected positions to correlate with transcriptional features, such as gene expression levels [50] or promoter regions. We analyzed the distribution of the identified positions along the MPXV genome (Fig.3, Fig. S4) and found that APOBEC-signature substitutions, consistently detected by all three applied methods, were relatively uniformly distributed across the genome (Kolmogorov-Smirnov test: human, p=0.44; non-human primates, p=0.23). Then, we analyzed potential correlations between the positions and known gene categories classified according to gene expression timing. We used a well-established categorization of MPXV genes into early, intermediate and late transcription [51] but found no statistically significant difference in the density of positions with APOBEC-signature substitutions among these gene categories (Chi-squared test: human, χ2=3.3, p=0.5; non-human primates, χ2=4.09, p=0.39). Among other genomic features associated with transcription in MPXV, we examined potential correlations with the positions of transcription start sites (TSS), transcription end sites (TES), and promoters, but found no statistically significant associations (human: TSS, p=0.37; TES, p=0.31; promoters, p=0.13; non-human primates: TSS, p=0.28; TES, p=0.37; promoters, p=0.11). Thus, our analysis of the genomic distribution of APOBEC-signature substitutions relative to transcription-linked genomic features found no significant associations.

### Secondary structure at APOBEC-signature substitution sites lacks known APOBEC-associated hotspot patterns

Previous studies have reported that several members of the APOBEC family preferentially mutate TC motifs within specific secondary structure elements [25]. An elevated frequency of APOBEC-induced mutations was observed in cytosines located at the 3’ end of hairpin loops, which can form in single-stranded DNA or RNA [26]. To analyze whether the genomic positions with APOBEC-signature substitutions detected in our study are also associated with this secondary structure pattern, we applied the bioinformatics tool RNAsselem, which had been previously developed by our group [39]. We extended the functionality of RNAsselem, which was initially developed for analyzing experimentally known secondary structures of RNA viruses, to support predicted secondary structures. Using RNAsselem, we analyzed elements of the predicted secondary structure at positions with APOBEC-signature substitutions and compared them to predicted secondary structure patterns at all other positions in the viral genome. We found no association of APOBEC-signature substitution sites with hairpin loop structures, nor did we observe any statistically significant differences between secondary structures at APOBEC-signature positions and at other positions in the MPXV genome (Fig. 4a). It was recently reported that mutations observed during the 2022 MPXV outbreak are enriched in inverted repeat (IR) regions [52]. However, our analysis of APOBEC-signature substitution sites within hairpin structures – which inherently include IR regions – contradicts this finding. To clarify this discrepancy, we extended our RNA-seq dataset from human and non-human primate hosts by including two additional mutation sets obtained from DNA sequencing data presented in [53] (hereafter referred to as DNA-seq:human1–2), as well as two published MPXV mutation datasets from the 2022 outbreak [54,55]. We then applied the Palindrome Analyser tool used in [52]. Our results (Fig. 4b) showed no statistically significant enrichment of APOBEC-signature substitution sites in IR regions of the MPXV genome in either the RNA-seq or DNA-seq datasets (Chi-square test: RNA-seq:human1, p=0.45; RNA-seq:human2, p=0.16; RNA-seq:monkey1, p=0.5; RNA-seq:monkey2, p=0.36; DNA-seq:human1, p=0.62; DNA-seq:human2, p=0.05; Isidro et al. [54], p=0.5; O’Toole et al. [55], p=0.38). Moreover, we did not observe any trend suggesting such enrichment.

**Figure 4.**
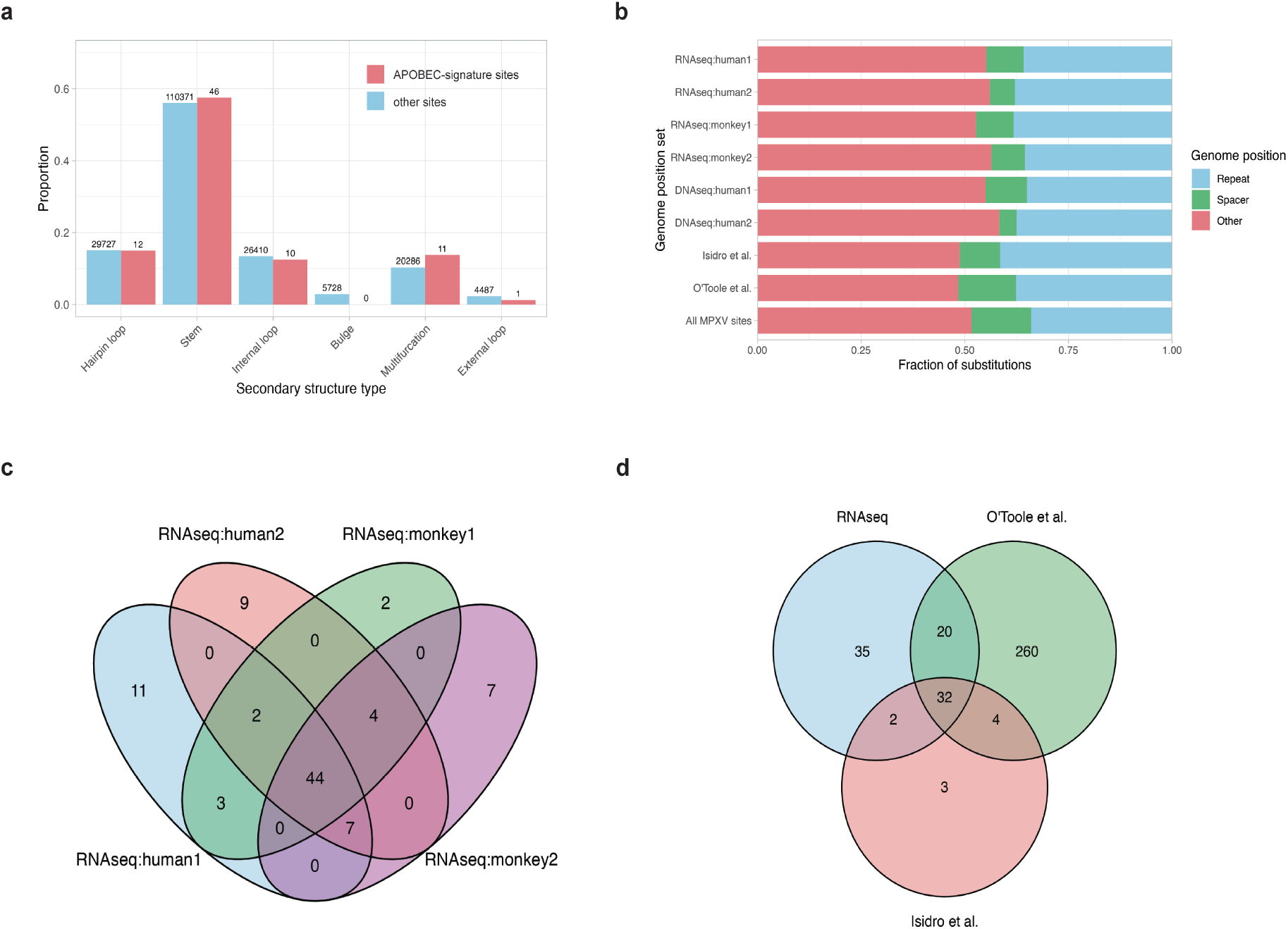
**(a)** Distribution of predicted secondary structure elements at the positions of observed substitutions in RNA-seq:human1-2 dataset and at all other genomic positions. No statistically significant enrichment of substitutions is observed in hairpin loops – structures previously associated with APOBEC mutational hotspots. **(b)** Distribution of APOBEC-signature substitutions identified in the datasets analyzed in this study and in previous works, mapped onto inverted repeat elements in the MPXV genome that are potentially capable of forming hairpin structures. No significant difference is observed between substitution sites and the rest of the genome. **(c)** Venn diagram showing the overlap of APOBEC-signature substitutions identified in human and monkey RNA-seq datasets analyzed in this study. **(d)** Venn diagram showing the intersection of APOBEC-signature substitutions detected across all RNA-seq and DNA-seq datasets analyzed in this study and mutations reported in MPXV strains from O’Toole et al. and Isidro et al.

### MPXV genome positions with APOBEC-signature substitutions significantly overlap across studies, reflecting pre-existing mutations rather than RNA editing hotspots

We also compared the lists of positions with APO-BEC-signature substitutions identified in the analyzed RNA-seq datasets, DNA sequencing studies and known MPXV mutation datasets [54,55]. A substantial overlap of substitution sites was observed across different datasets (Fig. 4c). Specifically, 53 of 80 sites (66%) overlapped between two human RNA-seq studies, 48 of 69 sites (70%) between two non-human primate studies, and 44 out of 89 sites (49%) were common across all four RNA-seq datasets. Comparison with the comprehensive catalog of MPXV mutations from the 2022 outbreak [54,55] revealed that 52 positions (58%) overlapped with those found in the RNAseq data (Fig. 4d). Notably, 38 of 41 positions (92%) reported in the earlier study by Isidro et al. [54] were also present in the union of sites detected in RNA-seq datasets and the O’Toole catalog [55].

To determine whether the observed substitutions occurred during the experiment or were already present, information on the specific MPXV strain used in each study was required. However, among the datasets considered, only the RNAseq:human2 project (PR-JNA906618) provided information on the specific MPXV strain used. For this project, we excluded from the list of detected substitutions known mutations of this strain, resulting in the exclusion of 54 out of 77 substitutions. For other projects, we excluded mutations listed in the comprehensive catalog of mutations observed in MPXV strains [55]. Thus, we obtained a conservative estimate of mutations that occurred during the experiments conducted in the studies considered and were not present in the strains used (Supplemental file 2). Although the number of these mutations is relatively small, there is a clear enrichment of APOBEC-signature mutations (22 out of total 49, p=2.19×10-8). The distribution of these mutations across the projects is not uniform: three out of six projects contain most of the mutations, while the remaining three have very few or none.

### Nucleotide context of substitutions suggests APOBEC3A or APOBEC3B as the likely enzymes responsible for APOBEC-signature mutagenesis

We further analyzed the trinucleotide context of APOBEC-signature substitutions identified in RNA-seq and DNA-seq studies (Fig. 5a). The most prevalent motif was TCG/CGA, accounting for 34% of all C→T/ G→A substitutions in the human RNA-seq data, 38% in the non-human primates RNA-seq, and 35% in the DNA-seq datasets. This was followed by the TCA/ TGA motif, which represented 30% of substitutions in the human RNA-seq, 28% in the non-human primates RNA-seq and 28% in DNA-seq samples. The TCT/AGA motif was the third most common motif, contributing 20% of substitutions in both human and non-human primates RNA-seq datasets and 21% in DNA-seq data. The fourth most frequent motif, TCC/ GGA, accounted for only ~7% of substitutions – substantially lower than other TC-containing motifs. Notably, this distribution differs from patterns observed in APOBEC-related cancer studies [49], where mutations in cytosines in the TCA motif are typically the most prevalent.

**Figure 5.**
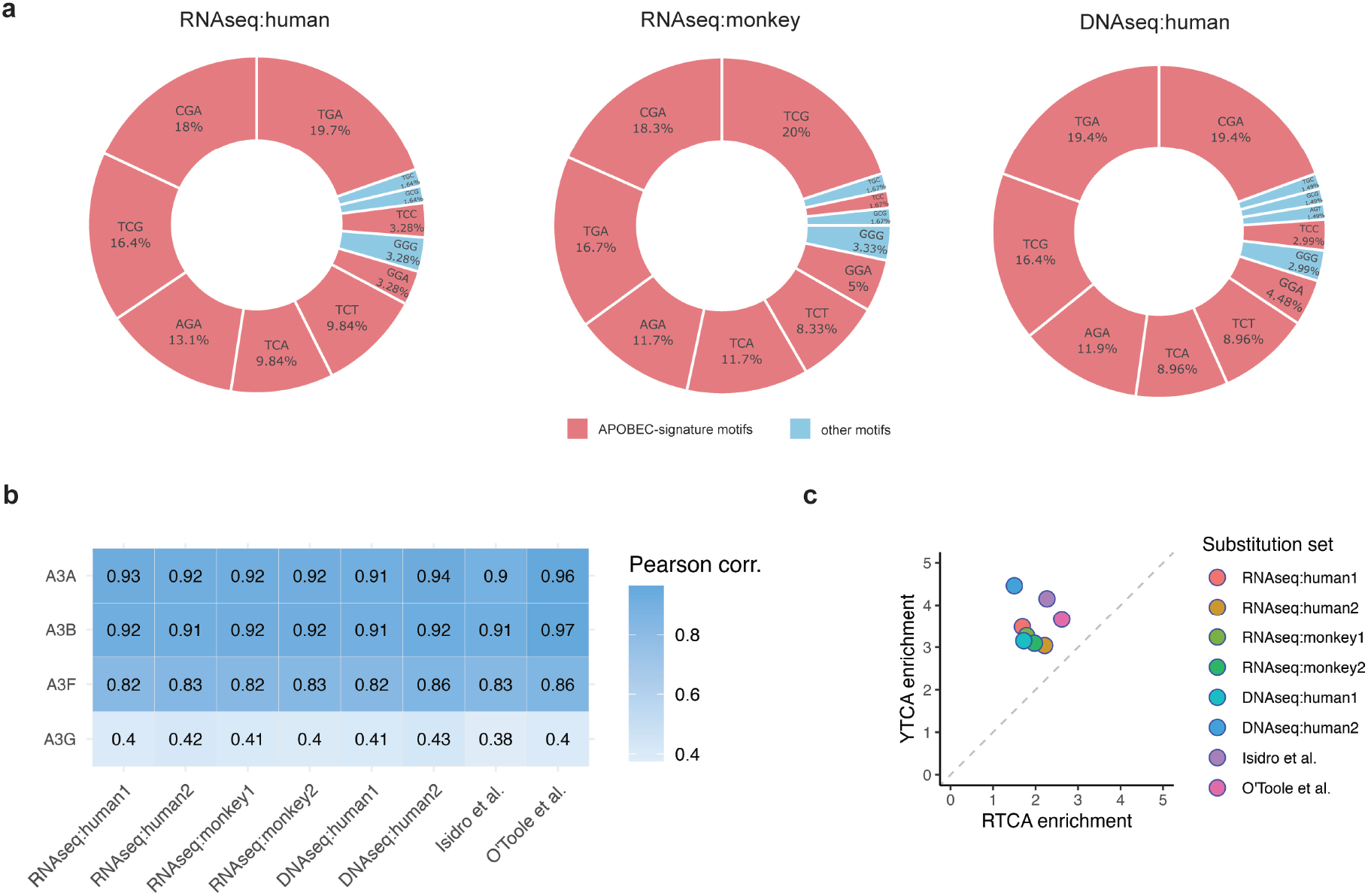
**(a)** Trinucleotide sequence context of C→T/G→A substitutions detected in RNA-seq and DNA-seq datasets from MPXV-infected human and monkey samples. **(b)** Correlation between the observed nucleotide context of substitutions and the known sequence preferences of APOBEC3 family enzymes. **(c)** Discrimination between the A3A- and A3B-associated mutational profiles using YTCA vs. RTCA motif analysis, where YTCA is preferentially targeted by A3A and RTCA by A3B.

We then analyzed the extended nucleotide context of APOBEC-signature substitutions to compare sequence profiles of these substitutions with known specificity models of APOBEC family members. First, we constructed positional weight matrices (PWMs) representing the sequence preferences of four APOBEC enzymes – A3A, A3B, A3G, and A3F – using available mutation data (see Methods). We then generated PWMs from the identified APOBEC-signature substitutions. The comparison of these matrices (Fig. 5b) revealed that the observed substitution profile most closely resembles the specificities of A3A and A3B. To differentiate between A3A and A3B, we conducted a YTCA vs. RTCA motif enrichment analysis [44], as these motifs are well-established signatures distinguishing these two enzymes. This analysis, applied across all datasets, consistently showed an enrichment of the YTCA motif (Fig. 5c), implicating A3A as the likely source of the observed mutations.

To complement this sequence-based inference, we additionally assessed the expression levels of APOBEC family members using RNA-seq data (Figure S5). In human infection datasets, we detected substantial expression of APOBEC3B, whereas APOBEC3A expression was essentially absent. In contrast, in the non-human primate datasets, APOBEC3A was prominently expressed, whereas APOBEC3B was also detectable but consistently at much lower levels than APOBEC3A. Thus, while at least one of the two candidate enzymes (APOBEC3A or APOBEC3B) was expressed in all infected samples, the expression patterns differed between human and monkey datasets. Taken together, although both the sequence context analysis and motif enrichment support APOBEC involvement, the heterogeneous expression profiles across datasets prevent a definitive assignment of the observed mutagenesis to either APOBEC3A or APOBEC3B alone. These results align with previous observations for MPXV mutations [27], but contradict an experimental study [56] suggesting APOBEC3F as the enzyme responsible for APOBEC-signature mutations in MPXV.

## Discussion

Interaction between viruses and host proteins play a crucial role in the viral life cycle. Viruses typically do not encode all proteins required for their life cycle, relying instead on host cellular proteins to perform essential functions. In contrast to interactions with recruited host proteins, viruses also engage in a constant battle with cellular defense factors — proteins whose primary function is to restrict viral replication. When a virus circulates within a host species for an extended period, it typically evolves mechanisms to evade cellular defenses. However, zoonotic transmission to a new host is usually associated with intensive interactions between the virus and the host’s antiviral proteins. This was partially observed during the SARS-CoV-2 pandemic and has become more pronounced in the recent monkeypox outbreak. Specifically, an accelerated accumulation of mutations exhibiting the signature of the APOBEC3 subfamily of viral restriction factors was observed in monkeypox virus strains circulating in humans during the 2022 outbreak [54,55]. Given that APOBEC3 enzymes can target both ssDNA and RNA [20], and that MPXV genomes and transcripts are localized in the cytoplasm during the viral life cycle [6], it remains an open question whether MPXV transcripts are also subject to APOBEC-mediated mutagenesis. In this study, we analyzed publicly available MPXV RNA-seq data to address this issue.

We considered RNA-seq data from two human and two non-human primate studies and developed a pipeline for detecting potential RNA editing events. Our analysis yielded consistent findings across all considered datasets, revealing a moderate enrichment of APOBEC-signature substitutions in MPXV transcripts. However, the majority of these substitutions occurred at high frequencies at specific positions within transcripts, leading us to conclude that either they are fixed mutations at the DNA level, or that they are RNA editing events occurring recurrently at specific hotspots. To distinguish between these two possibilities, we performed a detailed analysis of the identified APOBEC-signature substitutions. Firstly, as APOBEC3 catalyzes C→T editing in mRNA, we found that many G→A substitutions, which are complementary to C→T changes, remained even after normalization relative to the gene strand orientation. Secondly, we found that the observed substitutions were more frequently located at synonymous positions than at non-synonymous ones. Thirdly, we analyzed the distribution of these substitutions along the MPXV genome and found no correlation with the locations of genes expressed at different stages of the transcriptional program (early, intermediate, or late). Fourthly, secondary structure patterns previously associated with APOBEC mutational hotspots were not observed at the substitution sites. Finally, we compared the sets of identified APO-BEC-signature substitutions with known mutation sites in MPXV strains and found a substantial overlap. This overlap may, in part, reflect the presence of pre-existing mutations in the analyzed samples, as the specific viral strains used in the RNA-seq studies were not clearly specified. Taken together, these observations led us to conclude that the identified APOBEC-signature positions represent substitutions that occurred at the DNA level rather than RNA editing events.

We also investigated which member of the APOBEC family is most likely responsible for the observed mutagenesis. Given that members of the APOBEC family differ to some extent in their substrate specificity, we modelled the specificity via position-weight matrices using existing data for the A3A, A3B, A3F, and A3G enzymes. By comparing these models to the specificity profile derived from the observed data, we found consistently across all datasets that the sequence context of the substitutions best matches the specificity of A3A and A3B enzymes. Distinguishing between A3A and A3B specificity had been extensively considered in the context of cancer data [44], and here we applied a similar approach, identifying A3A as the likely mutator. However, expression analysis showed that the relative abundance of APOBEC3A and APOBEC3B varied across samples: in some datasets APOBEC3B was the predominantly expressed enzyme, in others APOBEC3A was more strongly expressed, and in a subset of samples both enzymes were co-expressed. This heterogeneity indicates that either A3A or A3B could, in principle, contribute to the observed APOBEC-signature mutations depending on the specific infection context. The known subcellular localization of A3A and A3B is more compatible with A3A activity, as A3B is predominantly localized in the nucleus, whereas A3A has been detected in both the nucleus and cytoplasm [57].

Although members of the APOBEC family are known to restrict a wide range of viruses, including both positive- and negative-sense single-strand RNA viruses, two distinct mechanisms may be involved: a deaminase-dependent pathway, which leads to C-to-T hypermutation, and a deaminase-independent pathway, in which APOBEC proteins inhibit viral replication through direct binding [58]. The deaminase-independent mechanism has been experimentally confirmed for both DNA and RNA viruses. Although it is more frequently associated with DNA viruses and retroviruses, evidence for its occurrence in RNA viruses [59,60], including reports of APOBEC-signature mutations in the positive-sense RNA virus SARS-CoV-2 [61,62].

Unlike RNA viruses, poxviruses replicate exclusively in the cytoplasm but generate single-stranded DNA intermediates during replication and repair, which represent plausible substrates for APOBEC3A and APOBEC3B. The APOBEC-associated mutational signature observed in contemporary MPXV genomes therefore likely reflects ongoing host-driven editing rather than replication errors intrinsic to the virus. Although the APOBEC signature does not directly determine clinical outcome, APOBEC-induced mutations may influence viral fitness, host adaptation, and lineage diversification, potentially affecting transmission dynamics during outbreaks. In this context, the enrichment of APOBEC-signature mutations in MPXV genomes provides insight into the interplay between host restriction mechanisms and viral evolution and may help explain the emergence of MPXV lineages with distinct epidemiological characteristics. Since APOBEC enzymes are known to mutate both viral DNA and RNA, the mechanisms preventing hypermutation of viral RNA transcripts remain poorly understood and require further research.

## Acknowledgements

This study was supported by Assignment FFRW-2024-0004 (to G.V.P., E.V.M., and R.K.A.) and, in part, by RSF grant #25-14-00491 (protein specificity modeling, to D.N.I). We thank Irina Ponomareva for designing the preprint layout.

## Data Availability

All scripts, source code, and intermediate data generated and used in this study are publicly available in the GitHub repository at: https://github.com/KazanovLab/APOBEC3-Mpox-RNA-seq. Raw sequencing data analyzed in this study are available from the corresponding NCBI BioProjects cited in the manuscript. Any additional information is available from the authors upon request.

