## Supplemental Figures and Tables for "Analysis of MPXV RNA-seq Data Reveals Lack of Evidence of APOBEC3-mediated RNA Editing"

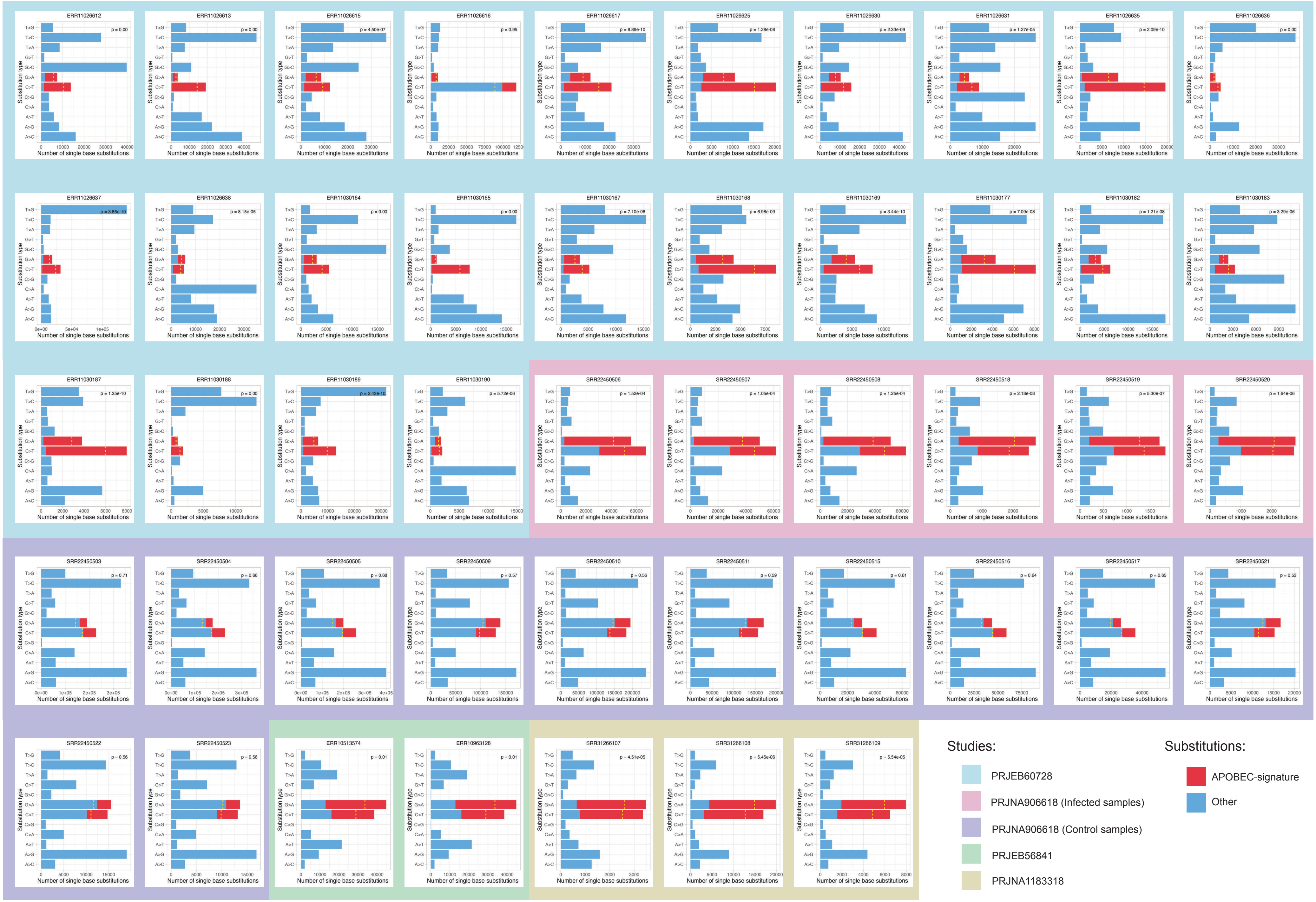


**Figure S1.** Distribution of observed substitutions in MPXV transcripts across substitution types. The dotted line marks the expected number of APOBEC-signature substitutions (C>T preceded by T and complementary G>A substitutions followed by A), as estimated from a beta-binomial model. The corresponding p-value indicates the significance of deviation from this expectation.

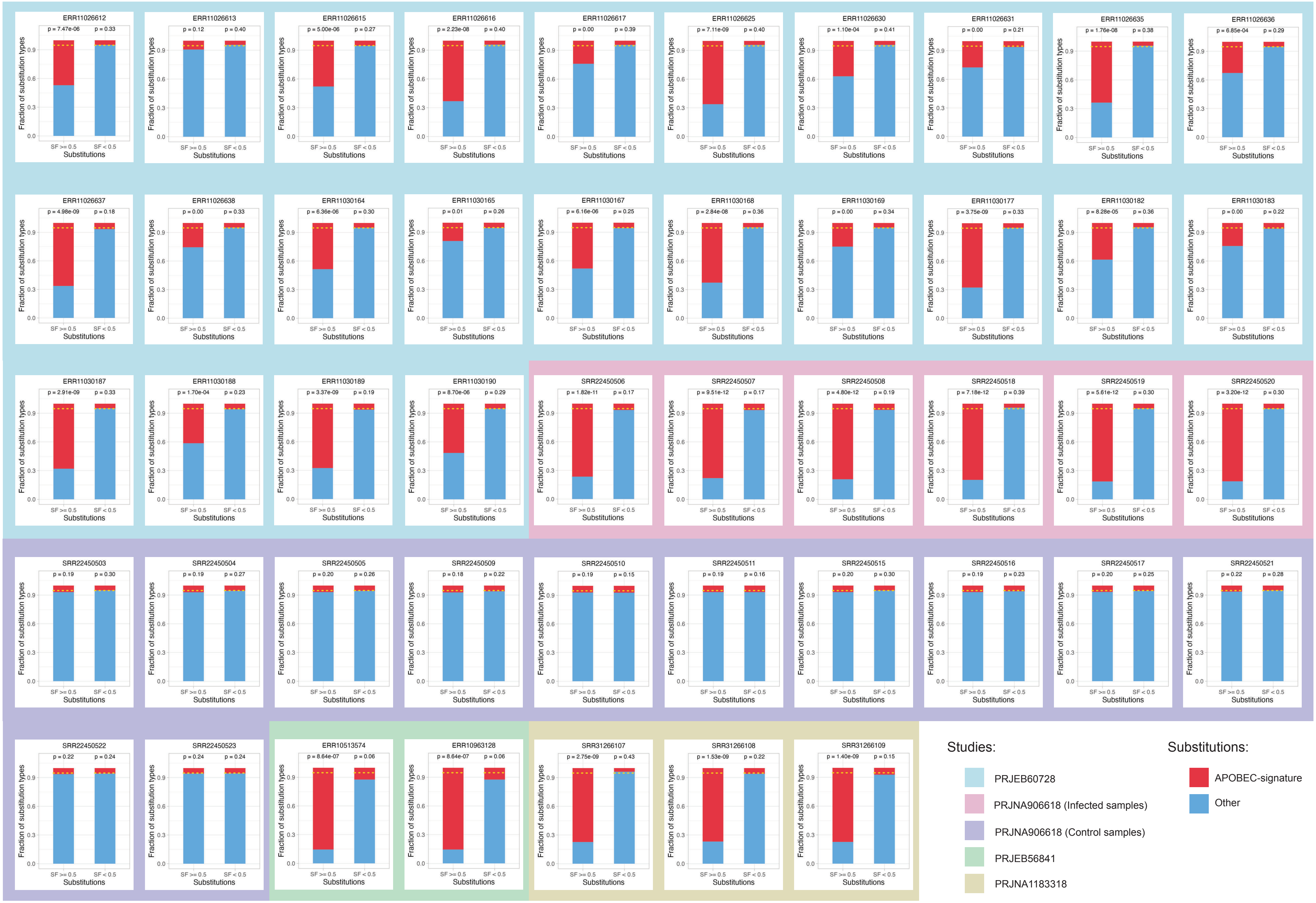


**Figure S2.** Enrichment analysis of APOBEC-signature substitutions across samples. Bars show the proportions of APOBEC-signature (red) and other (blue) substitutions within two substitution-frequency categories: high-frequency (SF ≥ 0.5) and low-frequency (SF < 0.5). Horizontal dashed lines represent the expected fraction of APOBEC-signature substitutions estimated from the beta-binomial model. P-values indicate the significance of deviation from this expectation for each sample.

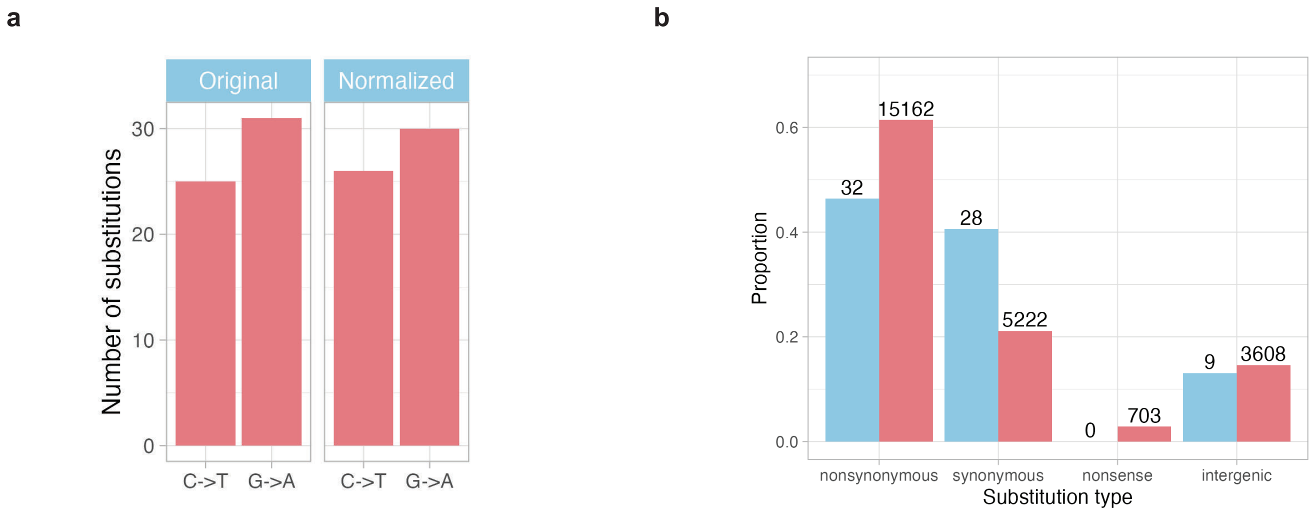


**Figure S3.** (a) The number of G→A substitutions identified in RNA-seq:monkey1-2 dataset remains elevated after normalization of C->T and G->A substitutions based on gene strand orientation. (b) Analysis of the effect of APOBEC-signature substitutions identified in RNA-seq:monkey1-2 dataset on the amino acid content encoded in MPXV transcripts reveals a prevalence of synonymous substitutions and a depletion of non-synonymous and nonsense substitutions.

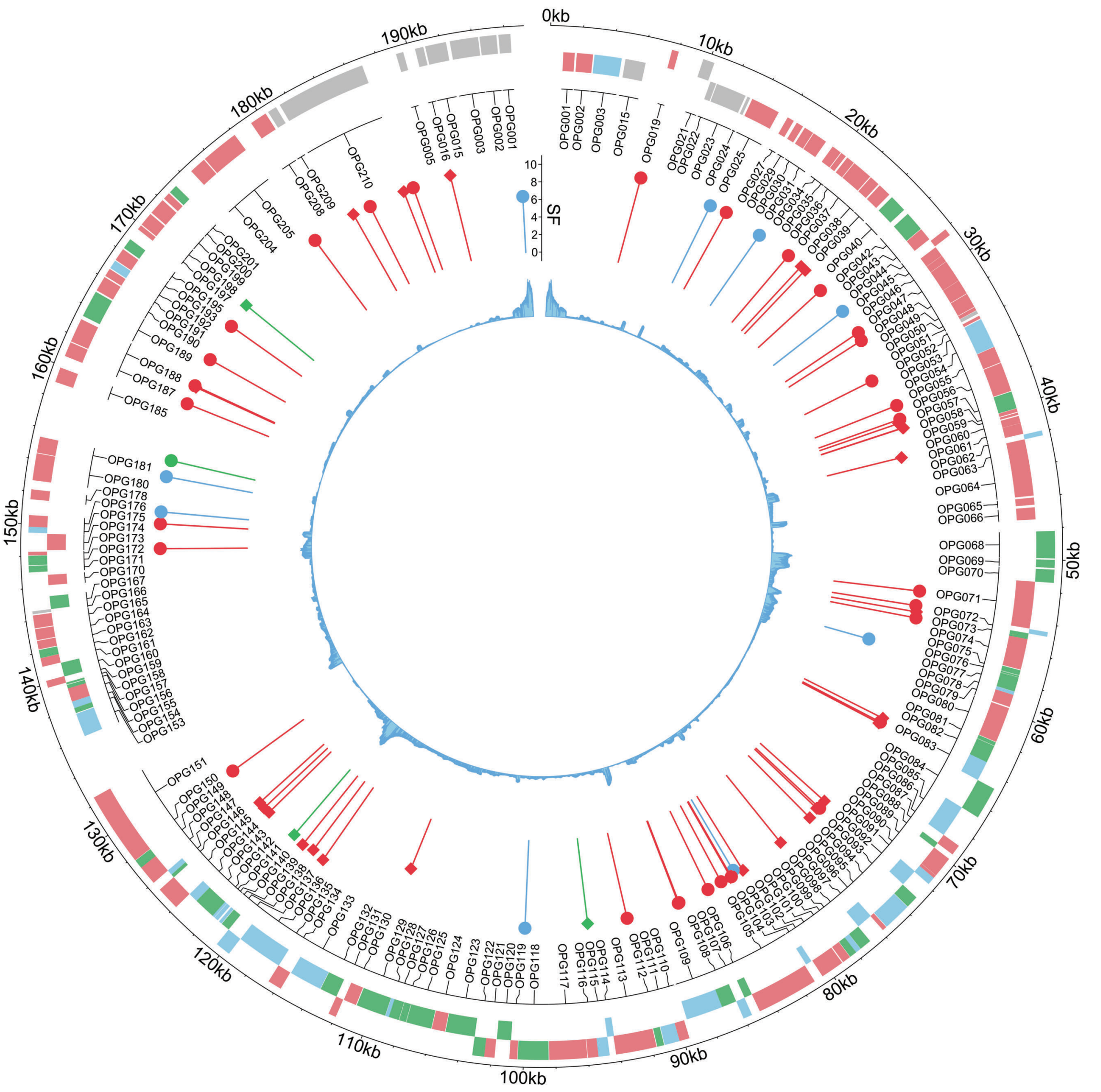


**Figure S4.** Genomic map of identified substitution positions in the MPXV genome from two monkey-derived projects. The outermost track shows annotated viral genes, color-coded by expression phase: early (green), middle (red), and late (blue). The next layer inward displays the positions of identified substitutions. The height of each pin reflects the frequency of the substitution. Pin color indicates substitution type: APOBEC-signature (red), extended APOBEC-signature (green), and other substitutions (blue). The pin head shape denotes strand orientation: a circle indicates a C->T substitution on the gene strand, while a square indicates the same substitution on the complementary strand. The innermost track displays the mean coverage across the genome.

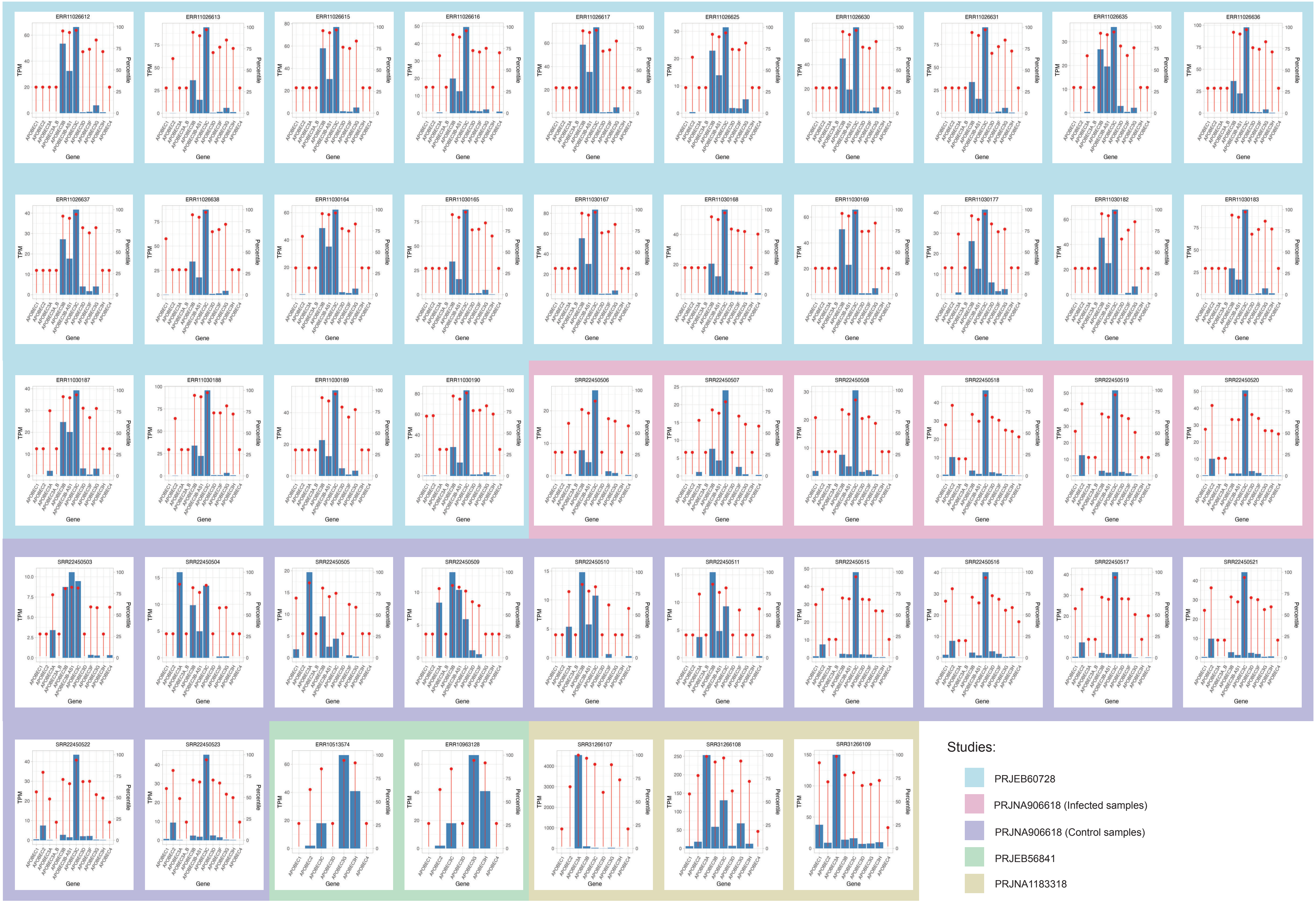


**Figure S5.** Each panel corresponds to a single sample and shows the expression of APOBEC genes (x-axis). Bars represent absolute transcript abundance in TPM (left y-axis). Red points denote the percentile position of each gene's TPM within the sample's full expression distribution (right y-axis), illustrating both absolute and relative expression patterns of APOBEC enzymes across the dataset.

| Accession | Project title | Organism | Data type | Number of samples |
|---|---|---|---|---|
| PRJEB60728 | Multi-omics characterization of the monkeypox virus infection | Homo sapiens | RNA-seq | 24 |
| PRJNA906618 | Virological characterization of the 2022 outbreak-causing monkeypox virus using human keratinocytes and colon organdies | Homo sapiens | RNA-seq | 18 (6 infected, 12 control) |
| PRJEB56841 | Long read, high-coverage transcriptome sequencing of Monkeypox virus and CV-1 host cell line on Oxford Nanopore Technologies flow cells in different time points of infection | Chlorocebus sabaeus | RNA-seq | 2 |
| PRJNA1183318 | A Penta-Component Mpox mRNA Vaccine Induced Protective Immunity in Naive and Simian Immunodeficiency Virus Infected Nonhuman Primates | Macaca fascicularis | RNA-seq | 3 |
| PRJNA845087 | Illumina whole-genome sequence of Monkeypox virus in a patient traveling from Canary Islands to France | Homo sapiens | DNA-seq | 4 |
| PRJNA981509 | Mpox reinfection in a vaccinated patient from Madrid (Spain) | Homo sapiens | DNA-seq | 2 |

Table S1. Summary of sequencing projects used in this study.

<table>
<tr><th>Method description</th><th>Command line</th></tr>
<tr><td>Alignment of long RNA reads to the host, viral, and hybrid reference genomes using Minimap2</td><td><code>$minimap -t $threads -a -x splice -Y -C5 --cs --MD<br>-un -G 10000 $ref $fastq | $samtools sort |<br>$samtools view --threads $threads --write-index -b<br>-o "${bam}"</code></td></tr>
<tr><td>Alignment of short RNA reads to the host, viral, hybrid reference genomes using STAR</td><td><code>"${star}" --genomeDir $ref_star --runThreadN<br>$threads --readFilesIn $fastq --readFilesCommand<br>"gunzip -c"  --outMultimapperOrder Random --<br>outSAMmapqUnique 60 --outSAMtype BAM<br>SortedByCoordinate --twopassMode Basic --<br>outFileNamePrefix "${sample_star_dir}" --outTmpKeep<br>All --outStd BAM_SortedByCoordinate --<br>outSAMunmapped Within KeepPairs --limitBAMsortRAM<br>1034634632 | $samtools view --threads $threads --<br>write-index -b -o "${bam}"</code></td></tr>
<tr><td>Alignment of short DNA reads to the host, viral, hybryd reference genomes using BWA</td><td><code>"${bwa2}" mem -R "@RG\tID:${sample}<br>\tPL:ILLUMINA\tLB:IDT\tSM:${sample}" "${ref}" -K<br>100000000 -M -Y $fastq -t 10  | $samtools sort --<br>write-index -m 500M -T "${sample_output_dir}/$<br>{sample}.sort" --threads 10 -o "${bam}"</code></td></tr>
<tr><td>Filtering of viral reads aligned to the hybrid genome from the BAM file</td><td><code>$samtools view --threads $threads -P $hybryd_bam --<br>write-index -b -o "${virus_bam_from_hybrid_bam}" “$<br>{virus_contig}”</code></td></tr>
<tr><td>Extraction of unmapped reads from the host-aligned BAM file</td><td><code>$samtools view --threads $threads -f 4 $host_bam --<br>write-index -b -o "${virus_bam_from_host_bam}"</code></td></tr>
<tr><td>Extraction of reads mapped to the viral genome from the BAM file</td><td><code>$samtools view --threads $threads -F 4 $virus_bam<br>--write-index -b -o "${virus_bam_from_virus_bam}"</code></td></tr>
<tr><td>Generation of FASTQ files from the filtered BAM file</td><td><code>$samtools collate --threads $threads  -u -O "$<br>{filtered_bam}" | $samtools fastq --threads<br>$threads -n -1 "${virus_fastq1_from_filtered_bam}"<br>-2 "${virus_fastq2_from_filtered_bam}" -0 "$<br>{virus_fastqS_from_filtered_bam}" -s "$<br>{virus_fastqO_from_filtered_bam}"</code></td></tr>
<tr><td>Variant calling using BCFtools with quality thresholds on mapping quality (MQ) and base quality (BQ)</td><td><code>$bcftools mpileup --threads $threads --min-MQ $MQ<br>--min-BQ $BQ --max-depth 1000000 --skip-indels  -f<br>$virus_ref -Ou "${realigned_bam}" | $bcftools norm<br>--threads $threads -m -snps --keep-sum AD --force |<br>grep -v "*" | $bcftools +fill-tags -- -t VAF |<br>$bcftools view --threads $threads --write-index -Oz<br>-o "${sample_output_dir}/${sample}_bcftools_MQ${MQ}<br>BQ${BQ}.vcf.gz"</code></td></tr>
</table>

Table S2. Command-line parameters used in the bioinformatic pipeline.
